# An anti-diuretic hormone receptor in the human disease vector, *Aedes aegypti*: identification, expression analysis and functional deorphanization

**DOI:** 10.1101/799833

**Authors:** Farwa Sajadi, Ali Uyuklu, Christine Paputsis, Aryan Lajevardi, Azizia Wahedi, Lindsay Taylor Ber, Andreea Matei, Jean-Paul V. Paluzzi

**Author notes:** **Corresponding Author:**, Lumbers Building, Room 221, Department of Biology, York University, 4700 Keele Street, Toronto, Ontario, M3J 1P3, Canada.

## Abstract

Insect CAPA neuropeptides, which are homologs of mammalian neuromedin U, have been described in various insect species and are known to influence ion and water balance by regulating the activity of the Malpighian ‘renal’ tubules (MTs). A number of diuretic hormones have been shown to increase primary fluid and ion secretion by the insect MTs and, in the adult female mosquito, a calcitonin-related peptide (DH_31_) also known as mosquito natriuretic peptide, increases sodium secretion at the expense of potassium to remove the excess salt load acquired upon blood-feeding. An endogenous mosquito anti-diuretic hormone was recently described, having inhibitory activity against select diuretic factors and being particularly potent against DH_31_-stimulated diuresis. In the present study, we have functionally deorphanized, both *in vitro* and *in vivo*, a mosquito anti-diuretic hormone receptor (*Aedae*ADHr). Expression analysis by quantitative PCR indicates the receptor is highly enriched in the MTs, and fluorescent *in situ* hybridization confirms expression within principal cells. Characterization using a heterologous system demonstrated the receptor was highly sensitive to mosquito CAPA peptides. In adult females, *Aedae*ADHr transcript knockdown using RNAi led to the abolishment of CAPA-peptide induced anti-diuretic control of DH_31_-stimulated MTs. The neuropeptidergic ligand is produced within a pair of neurosecretory cells in each of the six abdominal ganglia, whose axonal projections innervate the abdominal neurohaemal organs (known as the perivisceral organs), where these neurohormones are released into the open circulatory system of the insect. Furthermore, pharmacological inhibition of PKG/NOS signalling abolished the anti-diuretic activity of *Aedae*CAPA-1, which collectively confirms the role of cGMP/PKG/NOS in this anti-diuretic signalling pathway.

**Significance:** Insects are by far the most successful and abundant group of organisms on earth. As a result of their small size, insects have a relatively large surface area to volume ratio, raising the potential for rapid gain or loss of water, ions and other molecules including toxins – a phenomenon that applies to insects living in both aquatic and terrestrial environments. In common with many other organisms, hormones are key regulators of the excretory system in insects, and numerous factors control the clearance of excess water and ions (i.e. diuretics) or retention of these elements (i.e. anti-diuretics). Here we characterized an endogenous anti-diuretic hormone receptor in the human disease vector, *Aedes aegypti*, demonstrating its expression is highly enriched in the Malpighian ‘renal’ tubules and is necessary for eliciting anti-diuretic control of this key component of the mosquito excretory system.

## Introduction

Neuropeptides are central regulators of behaviours and control a plethora of physiological processes in all eukaryotic organisms. Insects, like many other animals, contain a comprehensive repertoire of neuropeptides along with their cognate receptors, which are essential for controlling complex biological phenomena including circadian rhythms, diapause, development, reproduction, pheromone biosynthesis, metabolism, circulation, stress as well as hydromineral balance^1–10^. Insects have a high surface area to volume ratio, which has implications for their ability to maintain levels of water and ions within a normal homeostatic range. In order to ensure their survival, most insects have a relatively ‘simple’ excretory system comprised of the Malpighian ‘renal’ tubules (MTs) and hindgut (ileum and rectum). The MTs produce the primary urine acting to clear the haemolymph of excess ions, metabolites and toxins while the hindgut generally functions in reabsorptive processes eliminating any unintentional loss of essential ions and amino acids^11,12^. The insect excretory system is under complex control, which may include direct innervation and regulation by neurotransmitters such as proctolin, as observed in the hindgut of many insects^13,14^. The excretory system in insects is also under the control by various circulating hormones^15,16^, which is the sole mechanism of extrinsic control in the non-innervated MTs, while endocrine factors may also influence the hindgut^4^.

The overwhelming majority of studies investigating regulators of the insect excretory system have focused on diuretic regulators of the MTs^17–25^, with only a few studies characterizing factors responsible for controlling reabsorptive processes across hindgut epithelia^12,26–30^. In addition, a few anti-diuretic factors that inhibit primary urine secretion by the insect MTs have also been reported^31–36^, acting to counter the activity of the diuretic hormones that increase ion and water secretion rates. We recently identified an endogenous anti-diuretic hormone in the disease-vector mosquito, *Aedes aegypti*, that strongly inhibits select diuretic factors including the mosquito natriuretic peptide (a calcitonin-related diuretic hormone)^37^, which is critical for the post-prandial sodium-rich diuresis that follows blood gorging by adult females^22^. Similarly, anti-diuretic activity of CAPA neuropeptides has been reported earlier in larval *A. aegypti* ^36^ as well as in other insects^31,38–42^, with signalling involving cGMP as a second messenger^31,37,40,42,43^. In addition to their clear anti-diuretic roles, CAPA peptides have also been linked to desiccation, where desiccation stress in *Drosophila melanogaster* leads to upregulation of *capa* mRNA, which is suggested to elevate CAPA levels in the CNS^44^. In many insects, CAPA peptides act through a conserved nitridergic signalling pathway leading to increased fluid secretion by MTs^24,44^. The mosquito anti-diuretic hormone is a member of the CAPA peptide family, which along with other insect PRXamide peptides, share homology to the vertebrate neuromedin U peptides^45^. CAPA neuropeptides are most abundant in specialized neurosecretory ventral abdominal (Va) neurons^46–49^ of the abdominal ganglia (or in the analogous neuromeres in insects with fused abdominal ganglia)^50,51^ and stored within abdominal perivisceral organs ^52–55^, which are major neurohaemal organs facilitating neurohormone release into circulation for delivery to target organs expressing receptors.

In the present study, we utilized a combination of molecular tools, heterologous functional assays, physiological bioassays and reverse genetics techniques to identify and unravel the functional role of an anti-diuretic hormone receptor in the disease-vector mosquito, *A. aegypti*. Our data provides further evidence that mosquito CAPA neuropeptides, together with their cognate receptor identified herein, function in a neuroendocrine system halting the stimulatory activity of diuretic hormones that, if left unregulated, may compromise ion and water homeostasis in this important anthropophilic mosquito.

## Materials and Methods

### Animals and dissections

Various stages of *A. aegypti* (Liverpool strain) were obtained from a laboratory colony maintained as described previously^56^. All mosquitoes were raised under a 12:12 light-dark cycle regime. Whole insects at each post-embryonic stage were used for examining developmental expression profiles and dissected tissues and organs were isolated from adults of each sex that were four-days post-eclosion. Adults were immobilized with brief exposure to carbon dioxide and then dissected to isolate individual organs using fine forceps (Fine Science Tools, North Vancouver, British Columbia, Canada) under nuclease-free Dulbecco’s phosphate-buffered saline (DPBS) at room temperature (RT).

### Immunohistochemistry

The dissected tissues/organs were fixed overnight at 4°C with 4% paraformaldehyde prepared in DPBS and were then washed several times with DPBS to remove fixative. The tissues were subsequently permeabilized in 4% Triton X-100, 10% normal sheep serum (NSS) and 2% bovine serum albumin (BSA) prepared in DPBS and incubated for 1 hour at RT on a rocking platform and then washed several times with DPBS to remove any traces of the permeabilization solution. The primary antibody was prepared using a custom affinity-purified rabbit polyclonal antibody (Genscript, Piscataway, NJ) produced against *Rhodnius prolixus* RhoprCAPA-2 (EGGFISFPRV-NH_2_; a kind gift from Prof. Ian Orchard, University of Toronto), which was diluted in 0.4% Triton X-100 containing 2% NSS and 2% BSA in DPBS. The stock antibody was diluted 1:1000 for stand-alone immunohistochemistry; however, when fluorescence *in situ* hybridization (FISH) preceded immunohistochemistry, the antibody was diluted 1:500 in the aforementioned solution. Tissues were incubated in the primary antibody solution for 48 hours at 4°C on a rocking platform, and no primary controls were incubated in the same solution of 0.4% Triton X-100 containing 2% BSA and 2% NSS in DPBS, but lacking primary antibody. After the primary antibody incubation, tissues were washed three times for one hour each with DPBS at RT. The secondary antibody solution was prepared using either FITC-conjugated sheep anti-rabbit immunoglobulin G (Jackson ImmunoResearch Laboratories, West Grove, PA) or Alexa Fluor 488-conjugated cross-adsorbed goat anti-rabbit immunoglobulin G (Life Technologies, Burlington, ON) diluted 1:200 in DPBS containing 10% NSS. The tissues were incubated in the secondary antibody solution overnight at 4°C on a rocking platform and were then washed with DPBS several times at RT. Tissues were mounted in ProLong Diamond Antifade Mountant containing DAPI (Molecular Probes, Eugene, OR) onto microscope slides and analyzed using a Lumen Dynamics XCite™ 120Q Nikon fluorescence microscope (Nikon, Mississauga, ON, Canada) or EVOS FL Auto Live-Cell Imaging System (Life Technologies, Burlington, ON).

### Determination of the complete cDNA of an *A. aegypti* anti-diuretic hormone receptor

The *Anopheles gambiae* CAPA receptor identified previously^57^ was used as a query for Megablast screening of the *A. aegypti* genomic scaffold database available locally on a lab computer running Geneious® 6.1.8 (Biomatters Ltd, Auckland, New Zealand) and the highest scoring hits (mapping within supercontig1.1) were assembled and predicted introns were excised. Using the Primer3 module in Geneious® 6.1.8, gene-specific oligonucleotides targeting this region (see Table S1) were designed to amplify this predicted partial fragment using Q5 High Fidelity DNA Polymerase (New England Biolabs, Whitby, On) with whole adult female *A. aegypti* cDNA as template. The PCR product was purified, A-tailed, cloned into pGEM-T vector (Promega, Madison, WI, USA) and nucleotide sequence was confirmed by Sanger sequencing (Center for Applied Genomics, Hospital for Sick Children, Toronto, ON). After successful validation of the cloned partial sequence, primers were designed (as described above) to perform 5′ and 3′ rapid amplification of cDNA ends (RACE)-PCR utilizing the Clontech SMARTer 5′/3′ RACE Kit (Takara BIO USA Inc, CA, USA) as recently described^58^. To facilitate cloning of amplicons, the linker sequence GATTACGCCAAGCTT, which overlaps with the pRACE vector provided in the RACE kit, was added to the 5′ ends of the gene-specific primers (Table 1). First-strand cDNA synthesis was prepared using 1μg total RNA from whole adult female mosquitoes using the 3′ CDS primer (provided in the kit) and a gene-specific reverse primer to generate template cDNA for 5′ RACE. Nested PCR reactions utilized gene-specific forward (3′ RACE) and reverse (5′ RACE) primers (see Table S1) and a universal primer mix (UPM) to amplify the complete cDNA encoding *A. aegypti* CAPAr with optimal cycling parameters determined empirically. Specifically, for both 5’ and 3′ RACE this included an initial denaturation at 94 °C for 1 min, followed by 40 cycles of 30 s at 94 °C, 30 s at 68 °C, and 3 min at 72 °C to amplify PCR products using SeqAmp DNA Polymerase. Following three rounds of nested PCR amplification using gene-specific primers, the amplicons were separated on a 1% agarose gel, extracted and cloned into the linearized pRACE vector. Plasmid DNA was isolated using a Monarch plasmid miniprep kit (New England Biolabs, Whitby, ON) and several clones were sent for sequencing for sequence validation. Finally, primers were designed at the 5′ and 3′ ends of the complete cDNA sequence (including UTRs) and used for final PCR amplification of the full receptor cDNA with Q5 High Fidelity DNA polymerase to confirm base pair accuracy.

### Heterologous receptor functional activation bioluminescence assay

The open reading frame of the cloned *A. aegypti* CAPAr was inserted into pcDNA3.1+ mammalian expression vector following procedures described previously^58–60^. Using a recombinant CHO-K1 cell line stably expressing aequorin^61^, *A. aegypti* CAPAr was transiently expressed following growth and transfection conditions as reported recently^58^. Cells were harvested for the functional assay at 48 hours post-transfection by detaching cells from the culture flask using 5mM EDTA in Dulbecco’s PBS (DPBS; Wisent Corp., St. Bruno, QC) and later cells were resuspended at a concentration of 10^6^-10^7^ cells/mL in assay media and incubated with coelenterazine *h* as described previously^60^. Prior to running the functional assay, cells were diluted 10-fold in assay media and left to incubate for one additional hour. Several endogenous as well as other insect neuropeptides representing a variety of neuropeptide families (see Table S2) were tested by preparing serial dilutions of each peptide in assay media. All peptides were commercially synthesized at a purity of >90% (Genscript, Piscataway, NJ) and 1mM stock solutions were prepared by dissolving 1mg of each peptide in water or DMSO as appropriate based on specific peptide characteristics. Recombinant CHO-K1 cells expressing the *A. aegypti* CAPAr were loaded into each well of multi-well plate using an automated injector module linked to a Synergy 2 Multi Mode Microplate Reader (BioTek, Winooski, VT) which measured kinetic luminescent response from each well for 20 sec immediately following cell loading onto the different peptides at various doses. Data was compiled in Microsoft Excel and analyzed in GraphPad Prism 8.0 (GraphPad Software, San Diego, CA).

### RNA probe template preparation

To obtain a template for synthesizing DIG-labelled RNA probes for use in FISH, a 373bp fragment of the *A. aegypti* CAPA partial mRNA (GenBank Accession: XM_001650839) previously described^50^ and a 743bp product of the anti-diuretic hormone receptor identified herein with primers designed (see Table S3) using the Primer3 plugin in Geneious® 6.1.8 (Biomatters Ltd., Auckland, New Zealand) were amplified using standard Taq DNA Polymerase (New England Biolabs, Whitby, ON) following manufacturer-recommended conditions. PCR products were column-purified with PureLink Quick PCR Purification Kit (Life Technologies, Burlington, ON) and amplified in a subsequent PCR reaction to generate cDNA products with incorporated T7 promoter sequence (see Table S1) to facilitate *in vitro* RNA synthesis of anti-sense or sense probes. The final purified PCR products for use as templates for RNA probe synthesis were quantified on a SYNERGY 2 Microplate reader (Biotek, Winooski, VT).

### Digoxigenin (DIG)-labelled RNA probe synthesis

PCR templates generated as described above (see Table S3) were used for *in vitro* transcription reactions using the HiScribe T7 RNA Synthesis Kit (New England Biolabs, Whitby, ON) following the recommended conditions when using modified nucleotides. Digoxigenin-labelled UTP was supplemented in a 35:65 ratio (DIG-UTP to standard UTP) either as a separate analog (digoxigenin-11-UTP) or in a pre-mixed 10x DIG-RNA labelling mix (Sigma-Aldrich, Oakville, ON). Template DNA was removed following treatment with RNase-free DNase I (New England Biolabs, Whitby, ON) and an aliquot of the synthesized RNA probes were then visually assessed using standard agarose gel electrophoresis and quantified on a SYNERGY 2 Microplate reader (Biotek, Winooski, VT).

### Fluorescence *in situ* Hybridization (FISH)

An optimized FISH procedure based on a protocol described previously for *R. prolixus*^62,63^ was utilized involving peroxidase-mediated tyramide signal amplification to localize cells expressing either the CAPA peptide mRNA or the anti-diuretic hormone receptor (CAPAr) mRNA. Tissues/organs were dissected under nuclease-free Dulbecco’s phosphate-buffered saline (DPBS; Wisent, St. Bruno, QC) and were immediately placed in microcentrifuge tubes containing freshly-prepared fixation solution (4% paraformaldehyde prepared in DPBS) and fixed for 1-2 hours at RT or overnight at 4°C on a rocker. Tissues/organs were subsequently washed five times with 0.1% Tween-20 in DPBS (PBT) and treated with 1% H_2_O_2_ (diluted in DPBS) for 10-30 minutes at RT to quench endogenous peroxidase activity. Tissues/organs were then incubated in 4% Triton X-100 (Sigma Aldrich, Oakville, ON) in PBT for 1 hour at RT to permeabilize the tissues and then washed with copious PBT. A secondary fixation of the tissues/organs was performed for 20 minutes in 4% paraformaldehyde in DPBS and then washed using PBT to remove all traces of fixative. The tissues/organs were then rinsed in a 1:1 mixture of PBT-RNA hybridization solution (50% formamide, 5x SSC, 0.1 mg/mL heparin, 0.1 mg/mL sonicated salmon sperm DNA and 0.1% Tween-20) which was then replaced with RT RNA hybridization that had been prepared earlier by denaturing in a boiling water bath for five minutes and subsequently cooled on ice for five minutes. The samples were then incubated at 56°C for 1-2 hours, which served as the pre-hybridization treatment. During the pre-hybridization incubation, labelled RNA probe (anti-sense for experimental or sense for control) was added to pre-boiled RNA hybridization solution (2-4ng/uL final concentration) and this mixture was heated at 80°C for 3 minutes to denature the single-stranded RNA probes and then cooled on ice for 5 minutes. The samples were then incubated overnight in this hybridization solution containing the DIG-labelled RNA probe at 56°C. The following day, samples were washed twice with fresh hybridization solution (minus probe) and subsequently with 3:1, 1:1 and 1:3 (vol/vol) mixtures of hybridization solution-PBT (all pre-warmed to 56°C). The tissues were subsequently washed with PBT pre-warmed to 56°C and in the final wash step were left to equilibrate to RT. Next, to reduce non-specific staining, samples were blocked with PBTB (DPBS, 0.1% Tween-20, 1% Molecular Probes block reagent; Invitrogen, Carlsbad, CA) for one hour. Tissues/organs were then incubated with a mouse anti-DIG biotin-conjugated antibody (Jackson ImmunoResearch Laboratories, West Grove, PA) diluted 1:400 and incubated for 1.5hrs at RT on a rocker in the dark. The antibody solution was then removed and tissues were subjected to several washes in PBTB over the course of one hour. Tissues/organs were then incubated with horseradish peroxidase-streptavidin conjugate (Molecular Probes, Eugene, OR) diluted 1:100 in PBTB for 1 hour and the tissues were once again washed with PBTB several times over the course of an hour. Finally, prior to treatment with tyramide solution for the signal amplification of the target mRNA transcripts, samples were washed twice with PBT and once with DPBS. Afterwards, a tyramide solution was prepared consisting of Alexa Fluor 568 (or Alexa Fluor 647) tyramide dye in amplification buffer containing 0.015% H_2_O_2_. After experimenting with various dilutions of the labeled tyramide, a 1:100 and 1:500 dilution of tyramide dye gave optimal results with minimal background staining for the ganglia and MTs, respectively. After the last DPBS wash was removed from the tissues/organs, the tyramide solution was added and the tissues were incubated in the dark for 1 hour on a rocker at RT. The tyramide solution was then removed and the samples were washed with DPBS several times over the course of an hour. The tissues/organs were stored in DPBS overnight at 4°C and then mounted on cover slips with mounting media comprised of DPBS with 50% glycerol containing 4 *μ*g/mL 4, 6-diamidino-2-phenylindole dihydrochloride (DAPI). For preparations involving transcript and neuropeptide co-detection in the nervous system, following the tyramide treatment, neural tissues were washed several times with DPBS and then incubated with primary antibody following the immunohistochemistry protocol described above. Tissues/organs were analyzed using a Lumen Dynamics XCite™ 120Q fluorescence microscope (Nikon, Mississauga, ON, Canada) or EVOS FL Auto Live-Cell Imaging System (Life Technologies, Burlington, ON).

### Synthesis of dsRNA for RNA interference and RT-qPCR

Double-stranded RNA (dsRNA) was synthesized and column-purified using the MEGAscript® RNAi Kit (Invitrogen, Carlsbad, CA) following the recommended protocol using primers for dsCAPAr synthesis (see Table S3) and primers as reported previously for dsARG^64^, which is an ampicillin resistance gene cloned from standard sequencing plasmid (pGEM T-Easy) that served as a negative control. A Nanoject Nanoliter Injector (Drummond Scientific, Broomall, PA) was used to inject one-day old female mosquitoes with 1μg (in ∼140nL) of either dsCAPAR or dsARG. After injection, mosquitoes were recovered in a photo period-, temperature- and humidity-controlled incubator. Total RNA was then isolated from four-day old whole female mosquitoes injected with dsCAPAr or dsARG using the Monarch Total RNA Miniprep Kit (New England Biolabs, Whitby, ON, Canada) and used as template (500ng) for cDNA synthesis using the iScript™ Reverse Transcription Supermix (Bio-Rad, Mississauga, ON, Canada) following recommended guidelines diluting cDNA ten-fold prior to quantitative RT-PCR. *Aedae*CAPAr and *Aedae*CAPA transcript levels were quantified using gene-specific primers that were positioned on different exons (see Table S3) and PowerUP™ SYBR® Green Master Mix (Applied Biosystems, Carlsbad, CA, United States) and measured on a StepOnePlus Real-Time PCR System (Applied Biosystems, Carlsbad, CA, United States) following conditions described previously^59^. A similar procedure for cDNA synthesis and transcript quantification as outlined above was followed for total RNA isolated from each post-embryonic developmental stage and tissues/organs dissected from adult stage mosquitoes. Relative expression levels were determined using the ΔΔCt method and were normalized to the geometric mean of *rp49* and *rps18* reference genes, which were previously characterized and determined as optimal endogenous controls^30^. Measurements were taken from three biological replicates, all of which included three technical replicates per reaction and a no-template negative control.

### Malpighian tubule fluid secretion assay

In order to determine fluid secretion rates, a modified Ramsay secretion assay^65^ was performed on isolated MTs of 3-6 day old adult female *A. aegypti,* as reported recently^37^. Tissue dissections were performed under physiological saline prepared as described previously^66^ diluted 1:1 with Schneider’s Insect Medium (Sigma-Aldrich, Oakville, ON). Individual MTs were removed and transferred to a Sylgard-lined Petri dish containing 20μl saline bathing droplets immersed in hydrated mineral oil to prevent evaporation. The proximal end of the MT was removed from the bathing saline and wrapped around a Minuten pin to allow for secretion measurements. Dosages of 25 nmol l^−1^ *Drome*DH_31_^22^ or 100 nmol l^−1^ 5-HT^67,68^ alone or in combination with 1 fmol l^−1^ *Aedae*CAPA-1 ^37^ were applied to the isolated MTs as previously described^37^. To investigate the effects of the pharmacological blockers, a nitric oxide synthase (NOS) inhibitor, N_ω_-Nitro-L-arginine methyl ester hydrochloride (L-NAME), and protein kinase G (PKG) inhibitor, KT5823, were used against 5-HT- and DH_31_-stimulated MTs. Dosages of 2 µmol l^−1^ L-NAME (manufacturer’s recommended dose) and 5 µmol l^−1^ KT5823^36^ were applied to the MTs. The inhibitors were treated in conjunction with 1 fmol l^−1^ *Aedae*CAPA-1 and/or 100 nmol l^−1^ cyclic guanosine monophosphate, 8 bromo-cGMP (cGMP)^37^ (Sigma-Aldrich, Oakville, ON, Canada). Unstimulated controls consisted of tubules bathed in physiological saline with no diuretic application. Following a 60-minute incubation period, the size of the secreted droplet was measured using an eyepiece micrometer and fluid secretion rate (FSR) was calculated as described previously^21^.

## Results

### Anti-diuretic hormone receptor identification and sequence analysis

The complete CAPA receptor in *A. aegypti* was identified and found to be 3461bp with an open reading frame of 2139bp encoding a receptor protein of 712 residues. The 5’ and 3’ untranslated regions were comprised of 899bp and 423bp, respectively (Figure S1A). The gene structure model revealed the cloned cDNA mapped to eleven exons spanning a genomic region of over 351Kb, with the start codon positioned within the third exon and the translation termination (stop) codon located in the eleventh exon, which is also contains the predicted polyadenylation signal at nucleotide position 3405-3410 (Figure S1B). The deduced protein sequence encodes a receptor protein that displays the prototypical features of rhodopsin receptor-like (family A) GPCRs ^69–71^, including the highly conserved tryptophan residue in the first extracellular loop involved in receptor trafficking, the D/E-R-Y/F motif at the border between the third transmembrane domain and second intracellular loop along with the NSxxNPxxY motif found within the seventh transmembrane domain (Figure S1A). Phylogenetic analysis using maximum likelihood methods revealed the deduced receptor protein sequence shares greatest evolutionary relationship with the orthologous CAPA receptor proteins identified or predicted in other dipterans organisms, including for example the fruit fly, non-biting midges, house fly, blow fly along with the more closely-related sister mosquito species (Figure S2).

### Functional ligand-receptor interaction heterologous assay

The endogenous peptidergic ligands for the cloned anti-diuretic hormone receptor were identified using a heterologous functional assay using CHO-K1 cells stably expressing a bioluminescent calcium sensor, aequorin^58,61^. The receptor was activated by all endogenously expressed peptides encoded by the CAPA gene in *A. aegypti* (Figure 1A), including two CAPA peptides (periviscerokinins) and a pyrokinin 1-related peptide. Notably however, the pyrokinin 1-related peptide displayed very poor activity compared to the two CAPA peptides, which were the most potent ligands with half maximal effective concentrations in the low nanomolar range (EC_50_ = 5.62-6.76 nM), whereas a significantly higher concentration of pyrokinin-1 was needed to achieve even low level CAPAr activation. Several other endogenous mosquito peptides as well as additional insect peptides belonging to distinct peptide families were tested and displayed no detectable activity over background levels of luminescence (Figure 1B). Controls where the CHO-K1-aeq cells were transfected with empty pcDNA3.1^+^ vector showed no detectable luminescence response (data not shown) to any of the peptides used in this study, confirming the calcium-based luminescence signal was a result of CAPA neuropeptide ligands activating the transiently expressed *A. aegypti* CAPA receptor.

**Figure 1.**
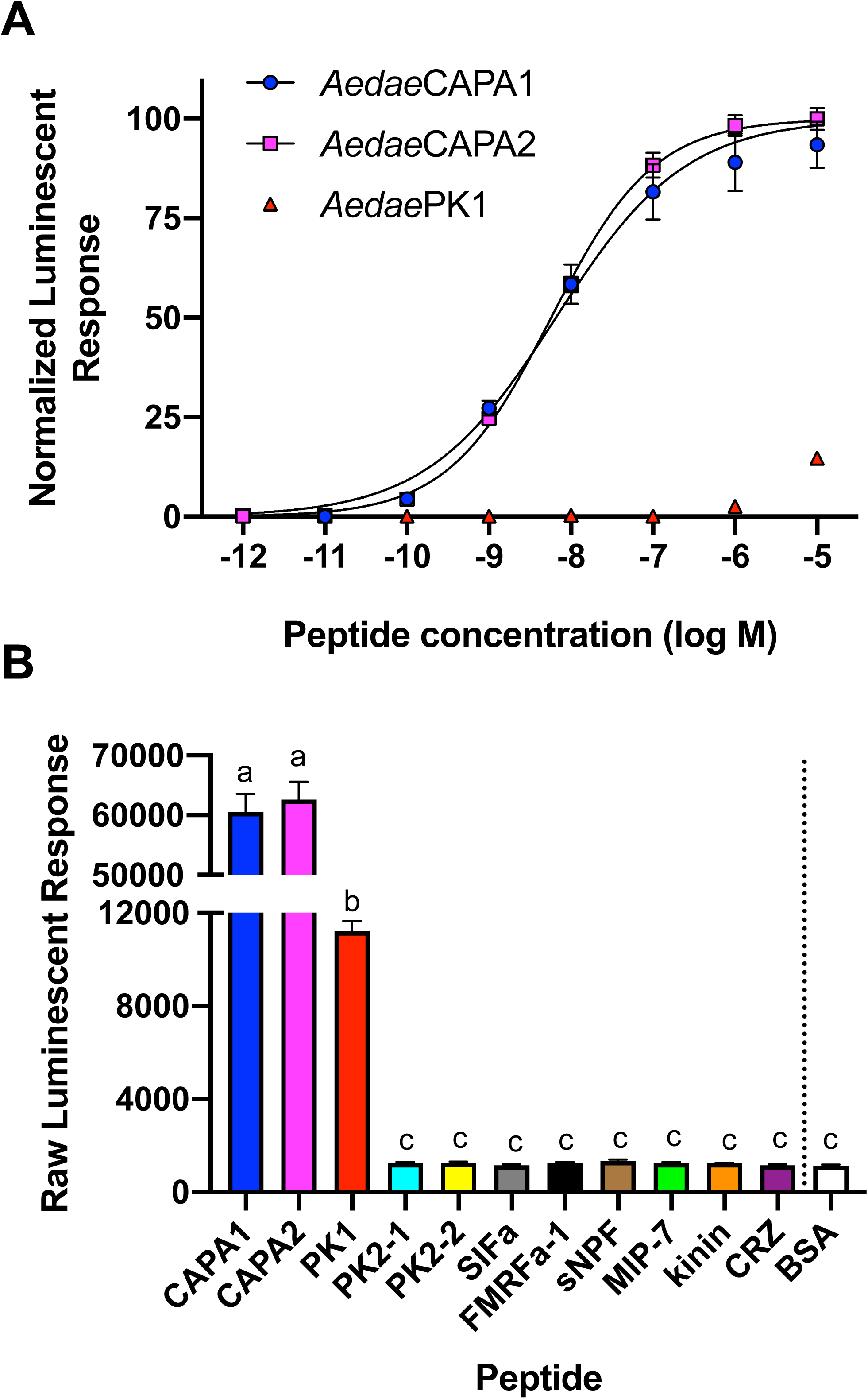
CAPA neuropeptide (anti-diuretic hormone) receptor (CAPAr) functional deorphanization using a heterologous assay. (A) Normalized dose-response curve demonstrating specificity of CAPAr functional activation by CAPA gene-derived neuropeptides. (B) Raw luminescent response following application of each CAPA gene-derived neuropeptide and representative neuropeptides belonging to several insect families, each tested at 10μM. For peptide sequence information and species origin, see Table S3. Only CAPA gene-derived neuropeptides resulted in a significant luminescent response relative to BSA control (vehicle). At this saturating dose, no difference in response was observed between the two endogenous CAPA neuropeptides, *Aedae*CAPA1 and *Aedae*CAPA2; *Aedae*PK1, demonstrated a significantly lower luminescent response (only ∼20% activity compared to either CAPA peptide), but nonetheless this response was significantly higher compared to all other tested peptides that were identical to background luminescent responses obtained with vehicle control (BSA). Different letters denote bars that are significantly different from one another as determined by one-way ANOVA and Tukey’s multiple comparison post-hoc test (p < 0.01). Data represent the mean ± standard error (n = 3).

### *CAPAr* transcript profile and cell-specific localization

We determined the developmental expression profile of the *A. aegypti* CAPA receptor (*CAPAr*) transcript in each post-embryonic developmental stage of the mosquito. Over the four larval stages and pupal stage of development, the *CAPAr* transcript level remained unchanged (Figure 2A); however, in adults, *CAPAr* transcript levels were significantly higher in adult male mosquitoes compared to adult female, pupal stage and first instar larval mosquitoes (Figure 2A). To confirm sites of biological action of the anti-diuretic hormones *in vivo*, we determined the *CAPAr* expression profile in adult *A. aegypti*, examining several tissues/organs in adult male and female mosquitoes. In males, *CAPAr* transcript was detected in reproductive tissues, head, carcass (i.e. the headless mosquito excluding the alimentary canal and reproductive tissues), midgut and low levels in the hindgut (Figure 2B). Enrichment of the *CAPAr* transcript was observed in the Malpighian ‘renal’ tubules (MTs) where expression was significantly enriched by ∼150-fold compared to all other tissues/organs examined (Figure 2B). A similar expression profile was observed in female mosquitoes with *CAPAr* transcript present in head, carcass, midgut and low levels detected in hindgut and reproductive organs. Similar to males, *CAPAr* was significantly enriched in the MTs of females relative to all other examined tissues/organs by nearly 150-fold (Figure 2B). Using fluorescent *in situ* hybridization, the *CAPAr* transcript was localized specifically to principal cells of the MTs and absent in stellate cells (Figure 2C). Specificity of *CAPAr* transcript localization was confirmed using sense control probe, with no signal detected in any cell type of the MTs (Figure 2D).

**Figure 2.**
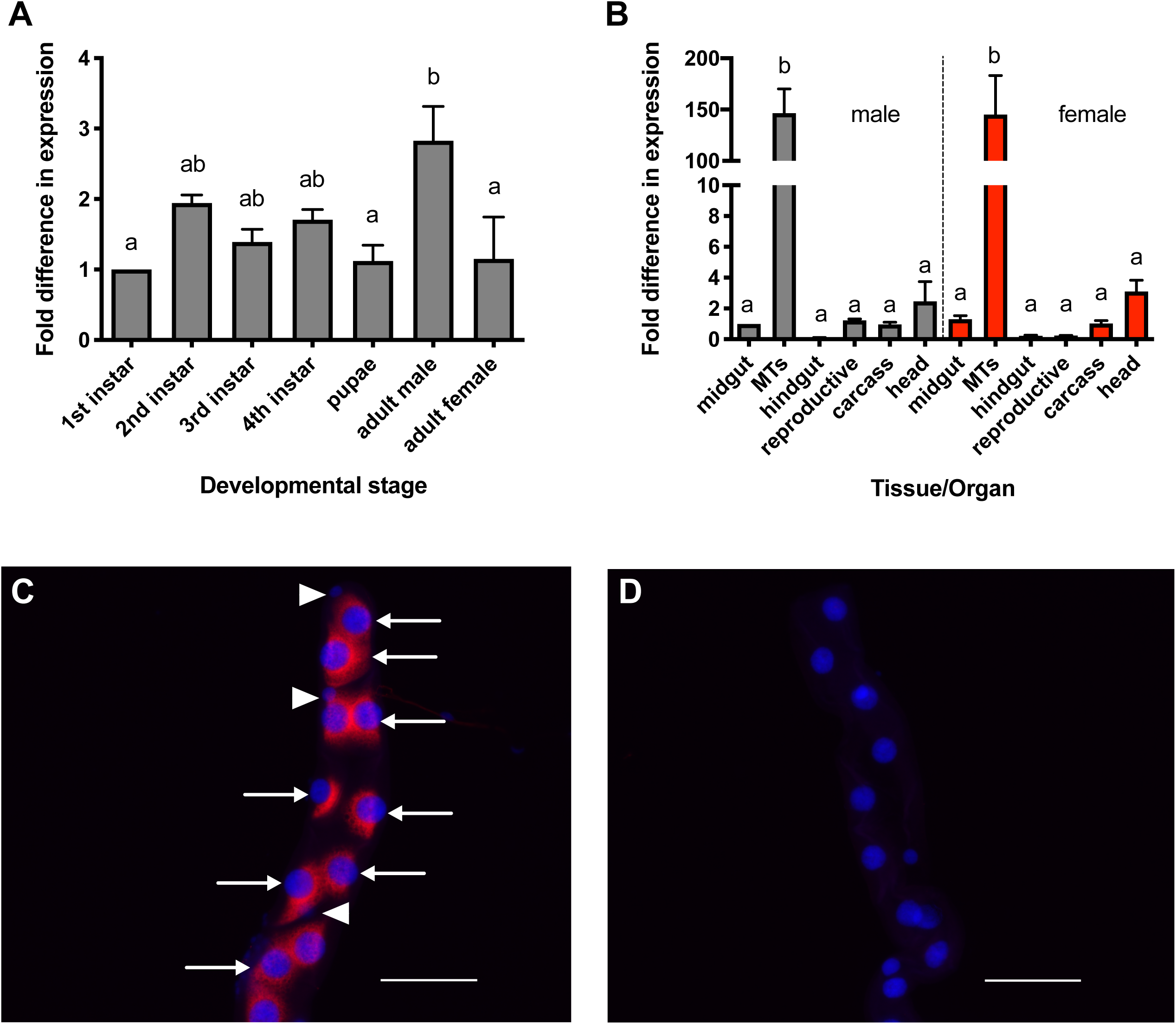
Expression analysis of *CAPAr* transcript in the mosquito, *A. aegypti*. (A) Ontogenic expression profile of *CAPAr* transcript over post-embryonic stages of the *A. aegypti* mosquito shown relative to transcript levels in 1^st^ instar larvae. (B) Spatial expression is analyzed in various tissues/organs from four-day old adult females, with transcript abundance shown relative to levels in the male midgut. (C) Cell-specific expression of *CAPAr* mRNA in principal cells (arrows) of MTs from adult female *A. aegypti* detected using an anti-sense probe, with no detection in the stellate cells (arrowheads). (D) No signal was detected in preparations hybridized with control *CAPAr* sense probe. All images acquired using identical microscope settings; scale bars in C-D are 100μm.

### *CAPA* transcript and mature neuropeptide immunolocalization within the abdominal ganglia

CAPA-like immunoreactivity was localized within all six of the abdominal ganglia, including the terminal ganglion. Specifically, each abdominal ganglion contains a pair of ventrally-localized strongly immunoreactive neurosecretory cells (Figure 3A). Axonal projections from these CAPA-like immunoreactive neurosecretory cells emanate dorsally and anteriorly within each ganglion, exiting via the median nerve (Figure 3B-C), with immunoreactive projections innervating the perivisceral organs (Figure 3D), which are the primary neurohaemal release sites in the ventral nerve cord facilitating neurohormone delivery into the insect haemolymph^72,73^. Validation that these immunoreactive neurosecretory cells in the abdominal ganglia were indeed CAPA-producing neurons was established by co-localization of CAPA transcript with CAPA-like immunoreactivity. Weakly staining CAPA-like immunoreactive cells were also observed in other regions of the central nervous system, including the brain, suboesophageal ganglion and thoracic ganglia (Figure S3); however, CAPA transcript was significantly enriched (∼140-fold) only within the abdominal ganglia but not in other regions of the nervous system (Figure S4). Within the abdominal ganglia, CAPA transcript co-localized within each pair of strongly staining CAPA-like immunoreactive neurosecretory cells (Figure 3F-H). Preparations treated with CAPA transcript sense probes did not detect any cells in the abdominal ganglia nor anywhere else in the central nervous system.

**Figure 3.**
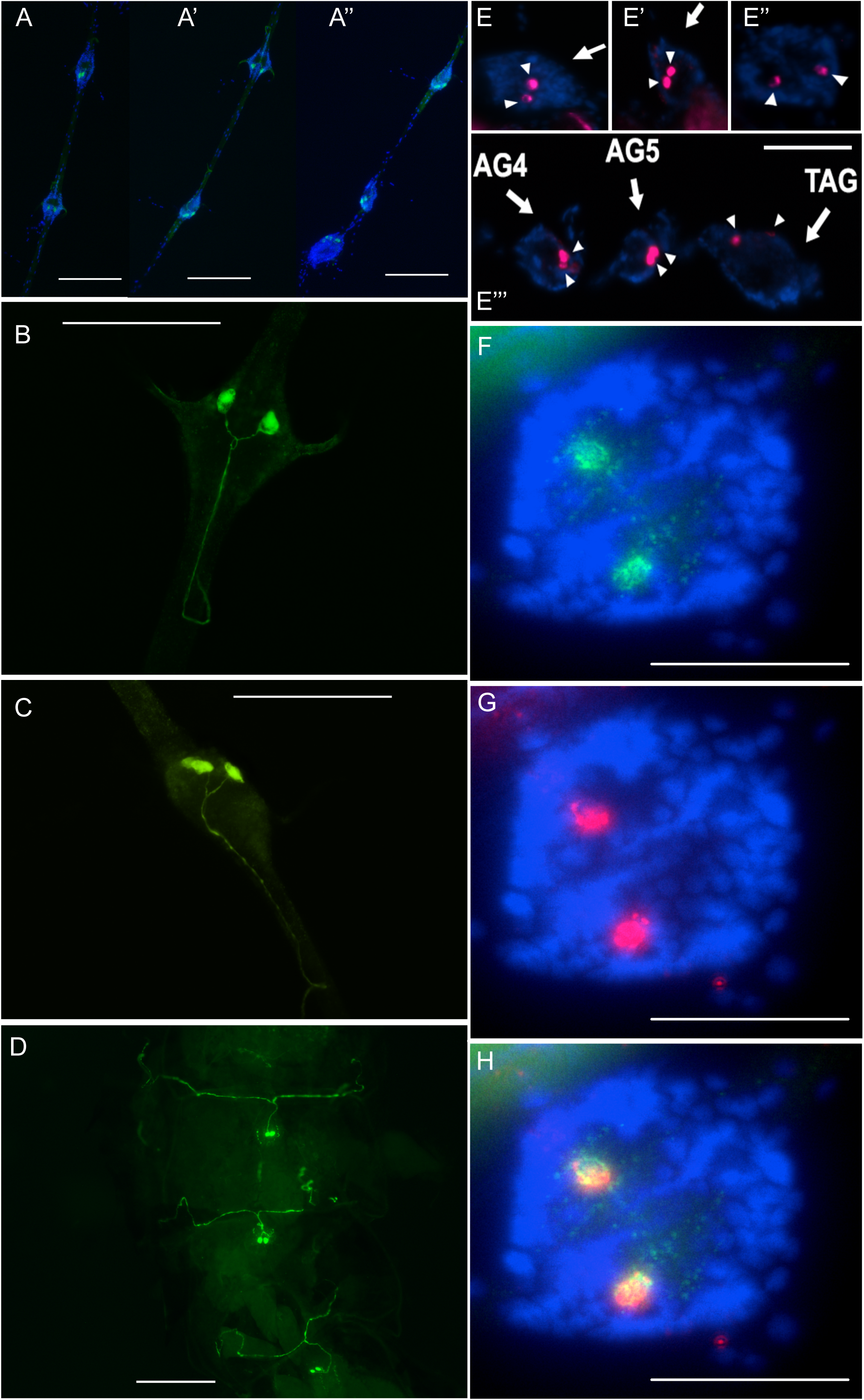
Mapping of anti-diuretic hormone in the abdominal ganglia of the central nervous system and associated neurohaemal organs in adult *A. aegypti*. (A) Immunohistochemical distribution of CAPA neuropeptides in the abdominal ganglia (AG); specifically, a pair of highly immunoreactive neurosecretory cells within AG1-2 (A), AG3-4 (A’) and AG4-6 (A’’). Higher magnification of AG3 (B) and AG4 (C) demonstrating CAPA immunoreactivity within large ventrally-positioned neurosecretory cells with axonal projections emanating dorsally within the ganglia and projecting anteriorly into the median nerve. (D) CAPA immunoreactivity in abdominal preparations with dorsal cuticle removed leaving the ventrally-localized AG within the abdominal segment showing immunoreactive processes innervating the abdominal neurohaemal (perivisceral) organs. CAPA transcript localization by fluorescent *in situ* hybridization revealing pairs of neurons within each AG including AG1 (E), AG2 (E’), AG3 (E’’) and AG4-5 and the sixth terminal abdominal ganglion (TAG; E’’’). Co-localization of CAPA immunoreactivity (F) and CAPA transcript (G) was verified in all abdominal ganglia with representative preparation in (H) showing transcript and immunoreactivity co-detection and overlap. Scale bars: 200µm (A & D), 100µm (B-C) and 50µm (E-H).

### *CAPAr* knockdown abolishes anti-diuretic hormone activity

To confirm that the anti-diuretic hormone action of the CAPA neuropeptides^36,37^ are mediated through this specific receptor expressed within the principal cells of the MTs, one day-old female *A. aegypti* were injected with ds*CAPAR* to knockdown *CAPAr* transcript levels. Relative to control mosquitoes injected with dsARG (gene cloned from standard plasmid vector encoding the ampicillin-resistance gene), *CAPAr* transcript was significantly reduced by ∼75% in four-day old females (Figure 4A). With significant knockdown verified in four-day old adult mosquito samples from the same experimental cohort, standard Ramsay assay was conducted as previously described^37^ on dsRNA-treated females to examine whether the anti-diuretic hormone activity of a CAPA anti-diuretic hormone was compromised. Our results confirmed that the CAPA neuropeptide, specifically *Aedae*CAPA-1, had no inhibitory activity against DH_31_-stimulated MTs in *CAPAr* knockdown females (Figure 4B). In contrast, *Aedae*CAPA-1 potently inhibited DH_31_-stimulated fluid secretion by MTs in dsARG-treated control female mosquitoes (Figure 4B).

**Figure 4.**
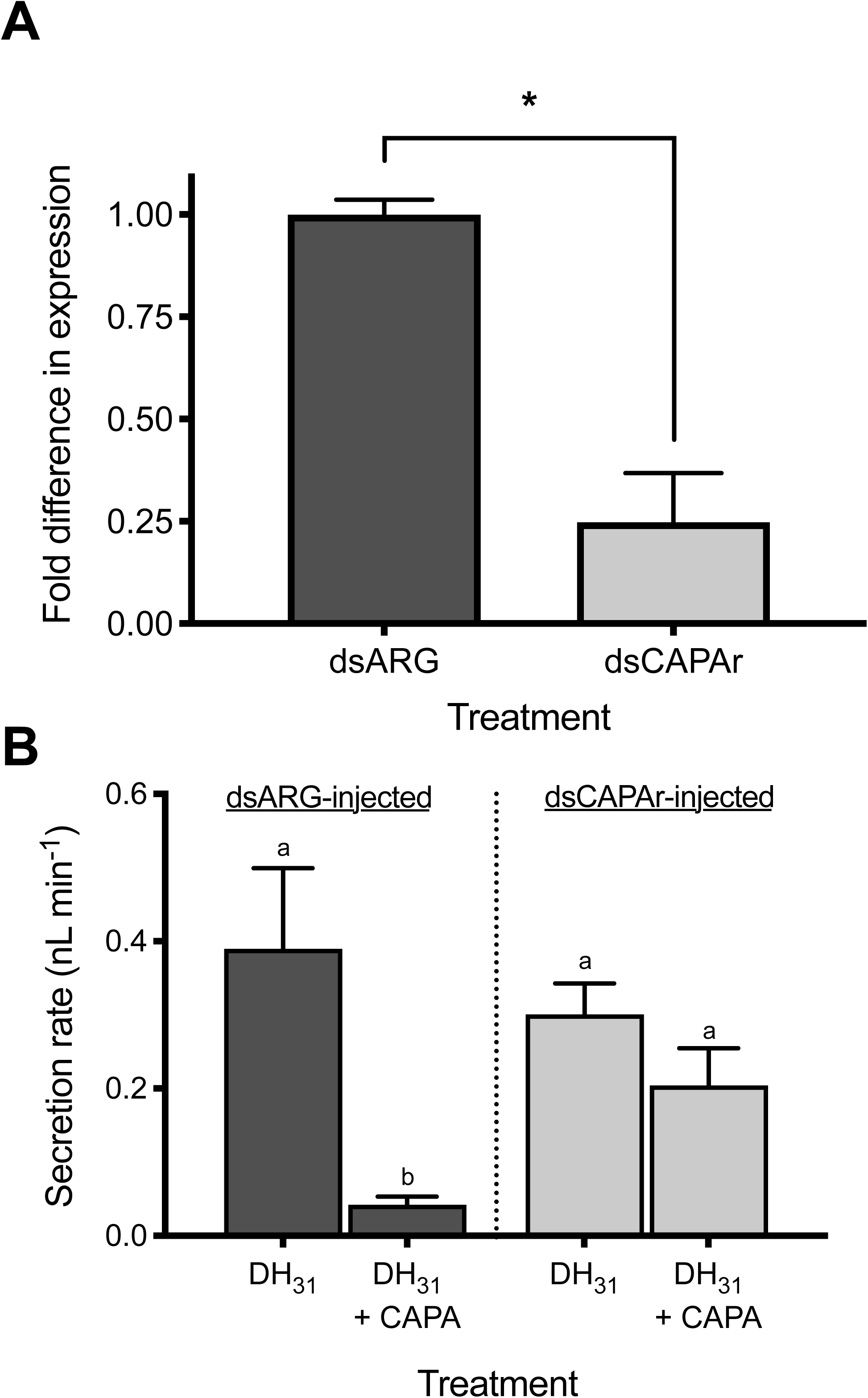
RNA interference (RNAi) of *CAPAr* abolishes anti-diuretic activity of CAPA neuropeptide on adult female *A. aegypti* MTs. (A) Verification of significant knockdown (>75%) of *CAPAr* transcript in MTs of four-day old adult female *A. aegypti* by RNAi achieved through injection of dsCAPAr on day one post-eclosion. (B) Functional consequences of *CAPAr* knockdown demonstrating loss of anti-diuretic hormone activity by *Aedae*CAPA-1 against *Drome*DH_31_-stimulated fluid secretion by MTs. In (A), knockdown of *CAPAr* transcript was analyzed by one-tailed t-test (* denotes significant knockdown, p < 0.01). In (B), fluid secretion rates by MTs presented as mean ± SEM and analyzed by one-way ANOVA and Tukey’s multiple comparison post-test, where different letters denote treatments that are significantly different (p < 0.05, n = 14-33).

### Effect of pharmacological blockade on the inhibition of fluid secretion

To further understand the anti-diuretic signalling pathway involving the CAPA neuroendocrine system, pharmacological blockers, including inhibitors of NOS (_L_-NAME) and PKG (KT5823), were tested against diuretic hormone-stimulated MTs alone and together with either *Aedae*CAPA-1 or cGMP. In DH_31_-stimulated MTs, _L_-NAME had no influence on the inhibitory effect of cGMP whereas the inhibitory effect of *Aedae*CAPA-1 was abolished (Figure 5A). In 5-HT-stimulated MTs, the results indicate that neither *Aedae*CAPA-1 nor cGMP inhibition are influenced by L-NAME (Figure 5B). Application of KT5823 abolished the inhibitory effect of both cGMP and *Aedae*CAPA-1 in DH_31_-stimulated MTs (Figure 6A). Similarly, the inhibitory activity of *Aedae*CAPA-1 and cGMP on 5-HT-stimulated tubules was abolished when treated with KT5823 (Figure 6B). Collectively, these results indicate that *Aedae*CAPA-1 inhibits select diuretic factors acting on the principal cells and involves NO and cGMP as a second messenger in DH_31_-stimulatd tubules, whereas cGMP, but not NO, is critical in the anti-diuretic activity of *Aedae*CAPA-1 on 5HT-stimulated MTs.

**Figure 5.**
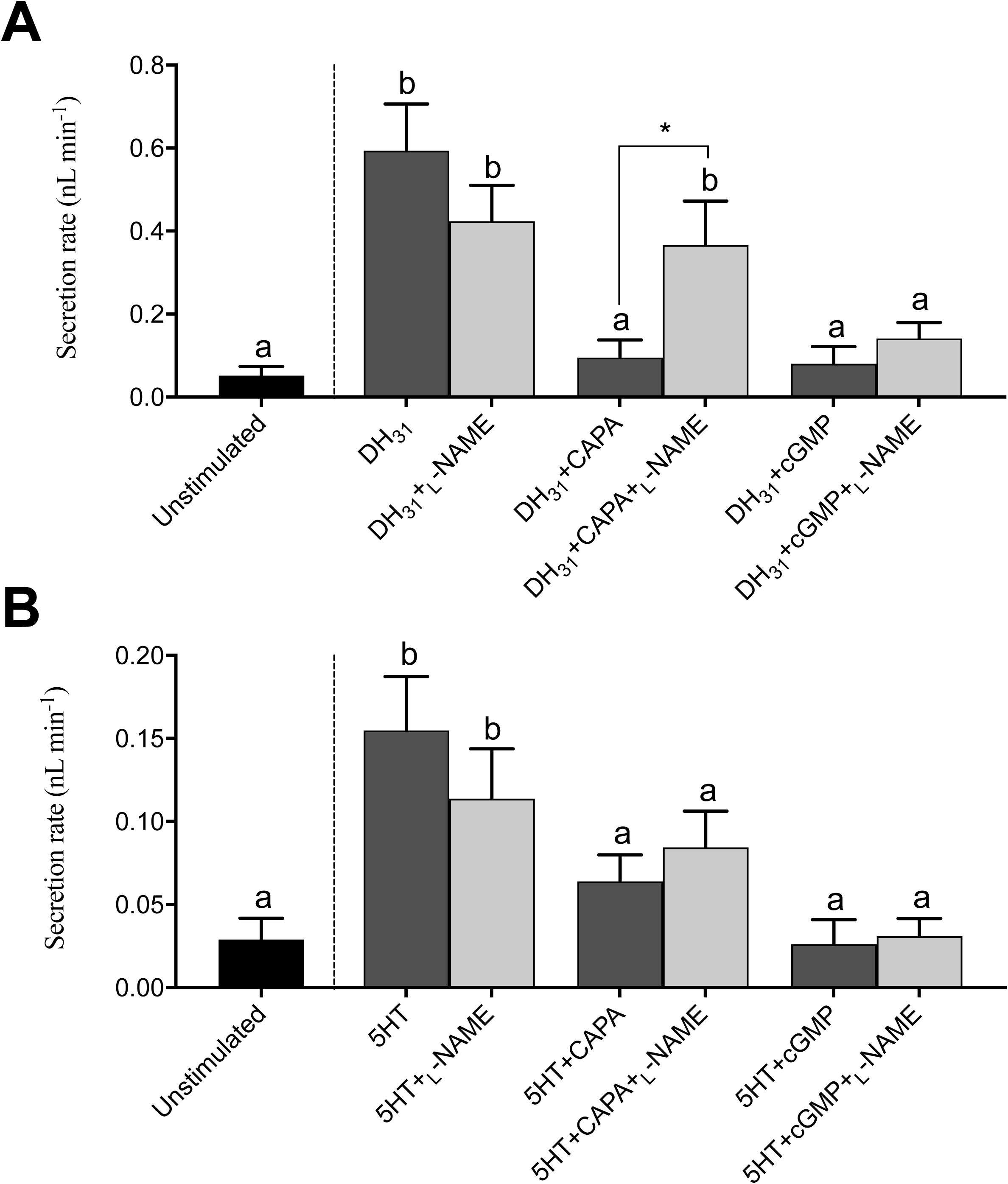
Effect of a nitric oxide synthase (NOS) inhibitor (_L_-NAME) on the anti-diuretic activity of *Aedae*CAPA-1 and cGMP in *Drome*DH_31_-stimulated *A. aegypti* MTs. The NOS inhibitor, _L_-NAME, was applied against (A) *Drome*DH_31_- and (B) 5HT-stimulated MTs alone or in the presence of either *Aedae*CAPA-1 or cGMP. Secretion rates are presented as mean ± SEM, n = 17-22. Columns that are significantly different from unstimulated controls are denoted with a distinct letter, as determined by a one-way ANOVA and Bonferroni post-test (p < 0.05).

**Figure 6.**
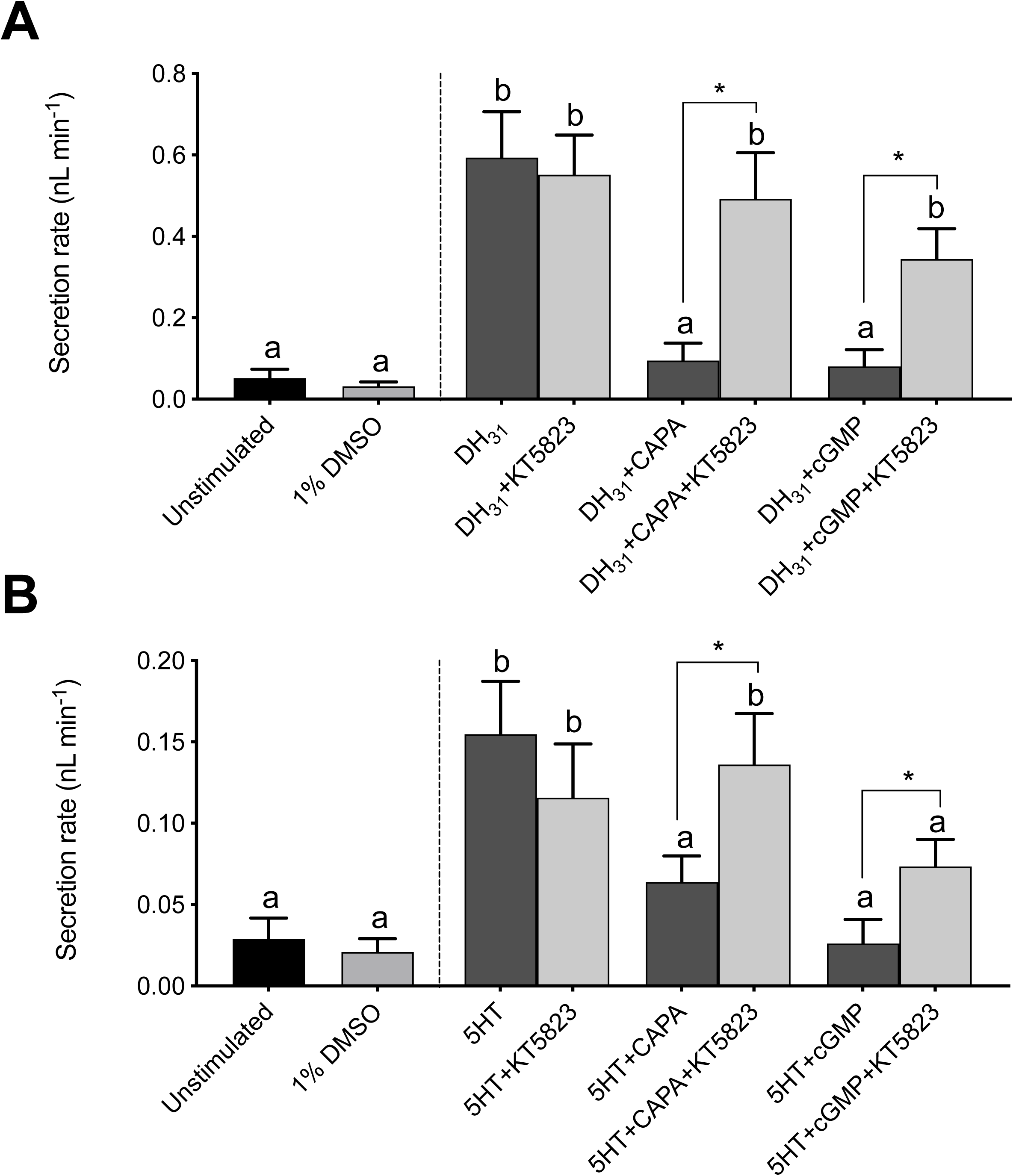
Effect of a protein kinase G (PKG) inhibitor (KT5823) on the anti-diuretic activity of *Aedae*CAPA-1 and cGMP in *Drome*DH_31_-stimulated *A. aegypti* MTs. The PKG inhibitor, KT5823, was applied against (A) *Drome*DH_31_- and (B) 5HT-stimulated MTs alone or in the presence of either *Aedae*CAPA-1 or cGMP. Secretion rates are presented as mean ± SEM, n = 16-25. Columns that are significantly different from unstimulated controls are denoted with a distinct letter, as determined by a one-way ANOVA and Bonferroni post-test (p < 0.05).

## Discussion

Like many animals, insects must regulate the ionic and osmotic levels of their internal environment to ensure homeostatic levels of water and electrolytes are maintained. This is critical not only for challenges linked to feeding, including the intake of too much or too little water and/or ions, but is also important for daily exchange of these elements with the environment through other routes such as waste elimination or water loss during respiration. The insect excretory system acts to maintain hydromineral balance of the haemolymph by either increasing the removal of water and/or ions in excess or the recycling of these same elements when in short supply. The insect Malpighian ‘renal’ tubules (MTs) play a key role as the organ responsible for primary urine production, which can then be modified by downstream elements of the excretory system such as the hindgut^4^. The MTs are the chief iono- and osmoregulatory organ and are under rigorous control by neuroendocrine factors, including both diuretic hormones (DH) and anti-diuretic hormones (ADH), which regulate transepithelial movement of ions and osmotically obliged water. These hormones consist of a variety of peptides as well as other neurochemicals produced by neurosecretory cells in the brain and ventral nerve cord^74,75^. Classically, DHs stimulate primary urine secretion by the MTs, whereas ADHs increase fluid reabsorption from the hindgut^15,76^. However, countless studies in diverse insect species have established that ADHs can also act on the MTs to reduce primary urine secretion^31, 36–38,40,77^. CAPA neuropeptides have been demonstrated to display potent anti-diuretic effects in a variety of insects^38,39,42,78^, including *A. aegypti* mosquitoes^36,37^, while they have been shown to function as DHs and ADHs in *D. melanogaster* ^40,79–81^.

The current study provides definitive evidence supporting the importance of this anti-diuretic hormone system in the disease vector mosquito, *A. aegypti*, by characterization and functional deorphanization of an anti-diuretic hormone receptor that is highly enriched in the MTs and demonstrates high selectivity for the mosquito CAPA neuropeptides. Previous studies have functionally deorphanized a number of CAPA receptor orthologs in other insects including dipterans^44,57,82,83^, lepidopterans^84^, coleopterans^85^, hemipterans^86^, as well as in the southern cattle tick^87^. Here, we have functionally validated the specific ligands of the elusive *A. aegypti* CAPA receptor demonstrating that two of the peptides encoded by the mosquito CAPA gene^50^, *Aedae*-CAPA1 and –CAPA2, potently activate this receptor leading to calcium signalling that elicits a bioluminescent response. While none of the other tested ligands representing multiple insect peptide families were active on the mosquito CAPA receptor, the third peptide encoded by the CAPA gene, *Aedae*-PK1, had low agonist activity with a potency of over five orders of magnitude lower compared to the canonical CAPA ligands. *Aedae*-PK1 is a member of the pyrokinin-1 family of peptides that contain the GXWFGPRL-NH_2_ (where normally X = V, M or L) consensus C-terminal sequence and recently a revised tryptopyrokinin nomenclature has been adopted to differentiate these neuropeptides from distinct pyrokinin families^88^. In agreement with our findings, a subset of previous studies on insect CAPA receptor orthologs have shown minor responsiveness to tryptopyrokinin ligands, with high doses eliciting low level CAPA receptor activation ^84–86^. Interestingly, this minor promiscuousness has not been observed for other dipteran CAPA receptors characterized previously^57,82,83^.

Members of the insect CAPA neuropeptide family are often also referred to as periviscerokinins due to their myotropic activity on visceral muscle and their source of release from the segmental abdominal neurohaemal organs known as perivisceral/perisympathetic organs^31,51,85^. Herein, we have immunolocalized CAPA neuropeptides within a pair of ventral neurosecretory cells within each of the six abdominal ganglia, whose axonal projections extend dorsally and anteriorly exiting each abdominal ganglion via the median nerve. CAPA immunoreactivity extends towards and is localized to the abdominal neurohaemal organs, the perivisceral organs, where these neuropeptides can be released into the haemolymph to elicit their neurohormonal actions on target sites expressing the CAPA receptor. The CAPA transcript was highly enriched within the abdominal ganglia of adult mosquitoes, confirming the transcript encoding the anti-diuretic hormone prepropeptide colocalized to these same neurosecretory cells. In support of these findings, peptidomic approaches using MALDI-TOF mass spectrometry have previously provided evidence for the presence of putative CAPA neuropeptides within isolated abdominal ganglia, including the terminal ganglion, from adult *A. aegypti*^50^. Collectively, these findings establish that the transcript and the mature peptide are present within the adult mosquito abdominal ganglia with the neurohormones being released into the insect circulatory system to act upon target tissues. Lastly, considering the low level CAPA transcript and immunoreactivity detected in other regions of the nervous system indicates that the abdominal ganglia, and their associated neurohaemal organs, are the primary source of the anti-diuretic hormone in adult *A. aegypti*. This also corroborates earlier peptidomic studies indicating the absence CAPA peptides, or differential processing of the CAPA precursor, in other regions of the nervous system aside from the abdominal ganglia where these neuropeptides are highly abundant^50,89,90^.

Having established the origin of the CAPA neuropeptide anti-diuretic hormones and their potent activity on the heterologously expressed CAPA receptor (CAPA-R), we next aimed to confirm the expression profile of the transcript encoding CAPA-R. Expression of *CAPAr* was observed in all post-embryonic ontogenic stages with significant enrichment in adult male mosquitoes, compared to females. Although the biological relevance of this differential expression remains unclear, this may relate to the sexual size dimorphism between adult male and female *A. aegypti*^91^, with the smaller males being inherently more susceptible to desiccation stress due to their higher surface area to volume ratio. In other insects, *CAPAr* transcript expression has been observed throughout most post-embryonic developmental stages^82,85,92^. The MTs are composed of two cell types forming a simple epithelium; large principal cells and thin stellate cells^74^. Principal cells facilitate the active transport of cations (Na^+^ and K^+^) into the lumen of the MTs from the haemolymph, while the stellate cells facilitate the transepithelial secretion of Cl^−^, the predominant inorganic anion^93^. In adult stages, expression analysis of *CAPAr* verified significant enrichment of this receptor in the MTs in both male and female mosquitoes. Furthermore, cell-specific expression mapping confirmed that the *CAPAr* transcript is restricted to the principal cells of the MTs and absent in the smaller stellate cells. In other insects, *CAPAr* transcript has been detected in various regions of the alimentary canal ^85,86,94^, including the principal cells of the MTs where this receptor is exclusively expressed in the fruit fly^44^. All in all, these observations are in line with physiological roles established for CAPA neuropeptides, which have been shown to modulate rates of fluid secretion by MTs in various insects ^36–37,41,62,96–97^. In dipterans, these effects are mediated via action on the principal cells acting through a second messenger cascade involving calcium, nitric oxide and cGMP signalling ^98,99^.

We next examined whether normal anti-diuretic hormone signalling, which requires the neuronally-derived CAPA peptide hormones activating their receptor expressed in the principal cells of the MTs, could be impeded by using RNA interference against the *CAPAr* transcript. One-day old mosquitoes were injected with *CAPAr*-targeted dsRNA resulting in knockdown at four-day old, where *CAPAr* transcript was significantly reduced. We examined whether *CAPAr* knockdown females retained sensitivity to CAPA peptides, which have been shown to inhibit fluid secretion by MTs by select diuretic hormones^37^. Indeed, *CAPAr* knockdown abolished the anti-diuretic activity of a CAPA neuropeptide against MTs stimulated with *Drome*DH_31_, an analog of mosquito natriuretic peptide. Collectively, through RNAi-mediated knockdown, these findings confirm that mosquito anti-diuretic hormones, which belong to the CAPA peptide family, are produced in pairs of neurosecretory cells in each of the abdominal ganglia whereby they are released through the neurohaemal organs and influence the MTs by acting on their receptor expressed within the principal cells of this organ. Further, the results confirm that sustained anti-diuretic hormone signalling, which requires the steady state expression of ligand and receptor, is necessary for facilitating the anti-diuretic control of the MTs.

In *D. melanogaster* and other dipterans, CAPA peptides have been shown to stimulate the nitric oxide (NO)/cGMP signalling pathway to induce diuresis^98^. When released, CAPA peptides bind to GPCRs found in principal (type I) cells of MTs, increasing Ca^2+^ levels in the cell through activation of L-type voltage gated calcium channels^100^. The influx of Ca^2+^ through these channels activates NOS, causing the production of NO, which subsequently activates guanylate cyclase to increase levels of cGMP in the MTs^44^. Ultimately, the activation of the NO/cGMP pathway stimulates the apical V-type H^+^-ATPase (proton pump), to increase fluid secretion in *D. melanogaster*. In the mosquito *A. aegypti*, CAPA peptides lead to activation of PKG, via elevated levels of cGMP^36^ and exogenous cGMP considerably inhibits fluid secretion rate^37^. Here, we sought to establish the roles of NO, cGMP and PKG on the anti-diuretic effects of CAPA peptides on adult mosquito MTs. Inhibitory doses of cGMP and a CAPA neuropeptide, namely *Aedae*CAPA-1, were treated with a NOS inhibitor, _L_-NAME, and a PKG inhibitor, KT5823. These investigations established that _L_-NAME did not alter the inhibitory effects of exogenous cGMP since this drug inhibits NOS, which is upstream of cGMP and, as a result, inhibition of DH_31_-stimulated secretion was unaffected. Contrastingly, *Aedae*CAPA-1 mediated inhibition of DH_31_-stimulated MTs was mitigated in the presence of _L_-NAME, reducing the anti-diuretic effects observed with *Aedae*CAPA-1. Comparatively, these findings are similar but are not identical to the effects of the PKG inhibitor, KT5823, which abolished the anti-diuretic activity of both *Aedae*CAPA-1 and cGMP, resulting in normal DH_31_-induced diuresis. Similar results were observed in 5-HT-stimulted MTs with one exception; *Aedae*CAPA-1 inhibition appeared to be independent of NOS since _L_-NAME had no influence on the anti-diuretic activity of *Aedae*CAPA-1 in 5-HT-stimulated MTs. Interestingly, the inhibition of both DH_31_- and 5-HT stimulated diuresis by *Aedae*CAPA-1 and cGMP were sensitive to the PKG inhibitor, KT5823, which indicates that while some differences in signalling associated with inhibition of different diuretic hormones may occur, these inhibitory pathways likely converge and involve cGMP activating protein kinase G. Taken together, the findings in this study provide definitive evidence that CAPA peptides are anti-diuretic hormones in the mosquito *A. aegypti*, which inhibit fluid secretion of adult mosquito MTs through a signalling cascade involving the NOS/cGMP/PKG pathway. Further studies are necessary in mosquitoes as well as other insects to elucidate the differential regulation by DHs and ADHs given ample data supporting that cGMP and related effectors can be both stimulatory^44,79,99,101^ and inhibitory^31,32, 36–37,43,81,102^ in their control on insect MTs. In conclusion, we have established an anti-diuretic hormone system in the adult mosquito *A. aegypti* providing evidence of a neural-renal axis whereby the neuropeptidergic anti-diuretic hormone is released by the abdominal segmental neurohaemal organs and subsequently targets their cognate receptor expressed within the principal cells of the MTs to counteract the activity of a subset of mosquito diuretic hormones. Fine-tuning of stimulatory and inhibitory hormones controlling the insect excretory system is of utmost importance to ensure overall organismal homeostasis to combat variable environmental conditions or feeding-related states that could perturb hydromineral balance if left unregulated.

## Supplementary file captions

**Figure S1.**
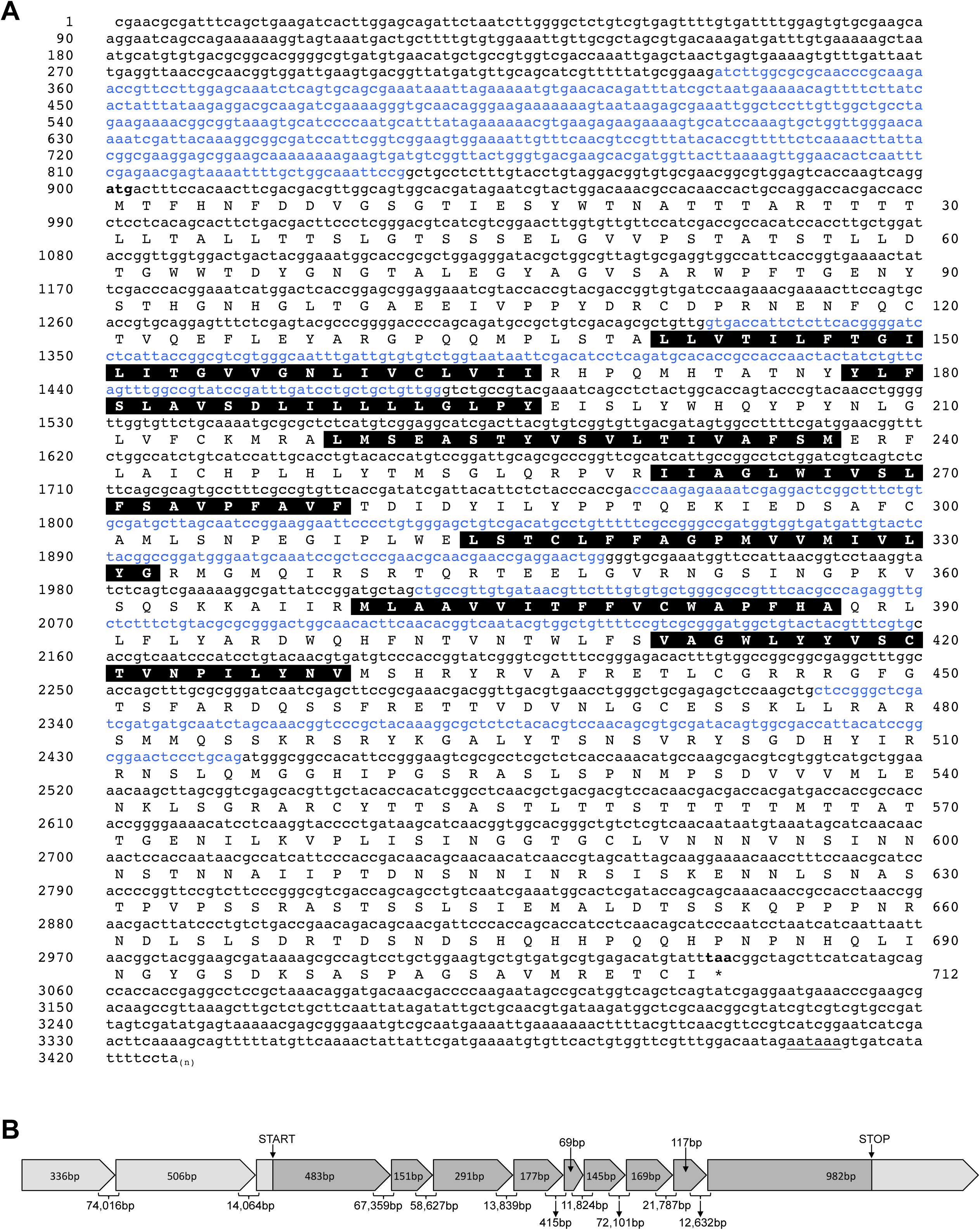
Sequence and gene structure of *A. aegypti* anti-diuretic hormone receptor. (A) The complete cDNA sequence (lowercase) and deduced protein sequence comprised of 712 amino acid residues (uppercase) along with predicted transmembrane domains (denoted by black highlighted residues) and other features as reported in the results text. Predictions of receptor features are described in the methods section. Nucleotides belonging to different exons are indicated by alternative blue/black font colour. Predicted polyadenylation signal is underlined in the 3’ untranslated region. (B) Exons with relative size to one another drawn to scale and denotes the open reading frame (in darker gray shading) beginning with the start codon in the third exon and stop codon within the eleventh exon. Intron sizes are predicted based on comparison of the deduced completed cDNA sequence with the *A. aegypti* genome scaffolds assessed on a local database using Geneious bioinformatics software (see methods for details). Predicted intron sizes range from as small as 415bp (between exons 6-7) and as large as 74,016bp (between exons 1-2) with the entire gene spanning a genomic region of >351kb.

**Figure S2.**
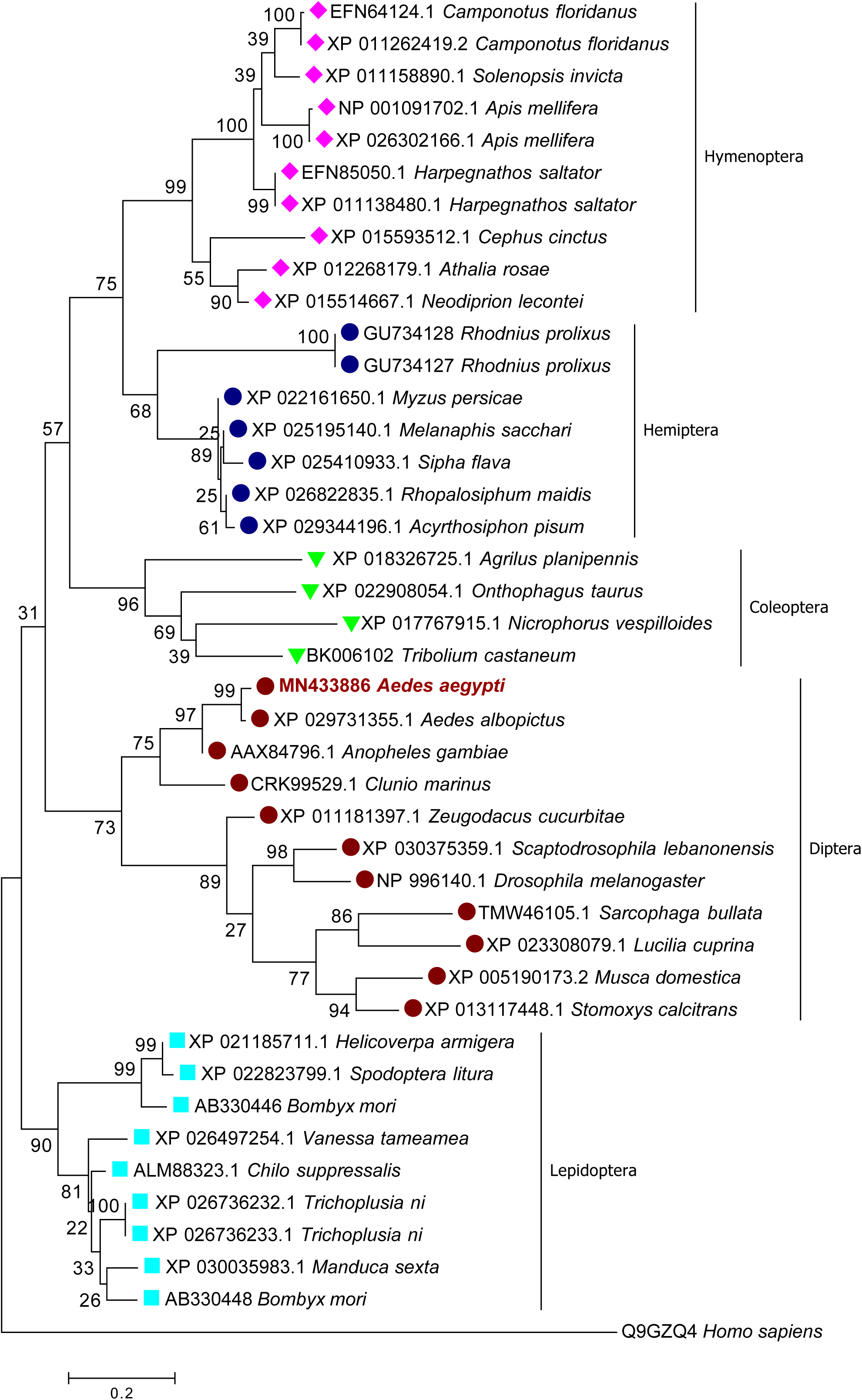
Molecular phylogenetic relationship of insect CAPA receptors inferred using the maximum likelihood method. Shown is the tree with the highest log likelihood with the numbers adjacent to the branches denoting the percentage of trees in which the associated taxa clustered together. A heuristic search was conducted to deduce an initial tree by applying Neighbor-Join and BioNJ algorithms to a matrix of pairwise distances estimated using a JTT model. Following this initial analysis, the topology with superior log likelihood value was selected automatically. Branch lengths are drawn to scale and denote the number of substitutions per site based on the final analysis involving 42 amino acid sequences and a total of 206 residue positions in the final data set with positions containing gaps and missing data removed. The human neuromedin U receptor 2 was included in the analysis and designated as the outgroup.

**Figure S3.**
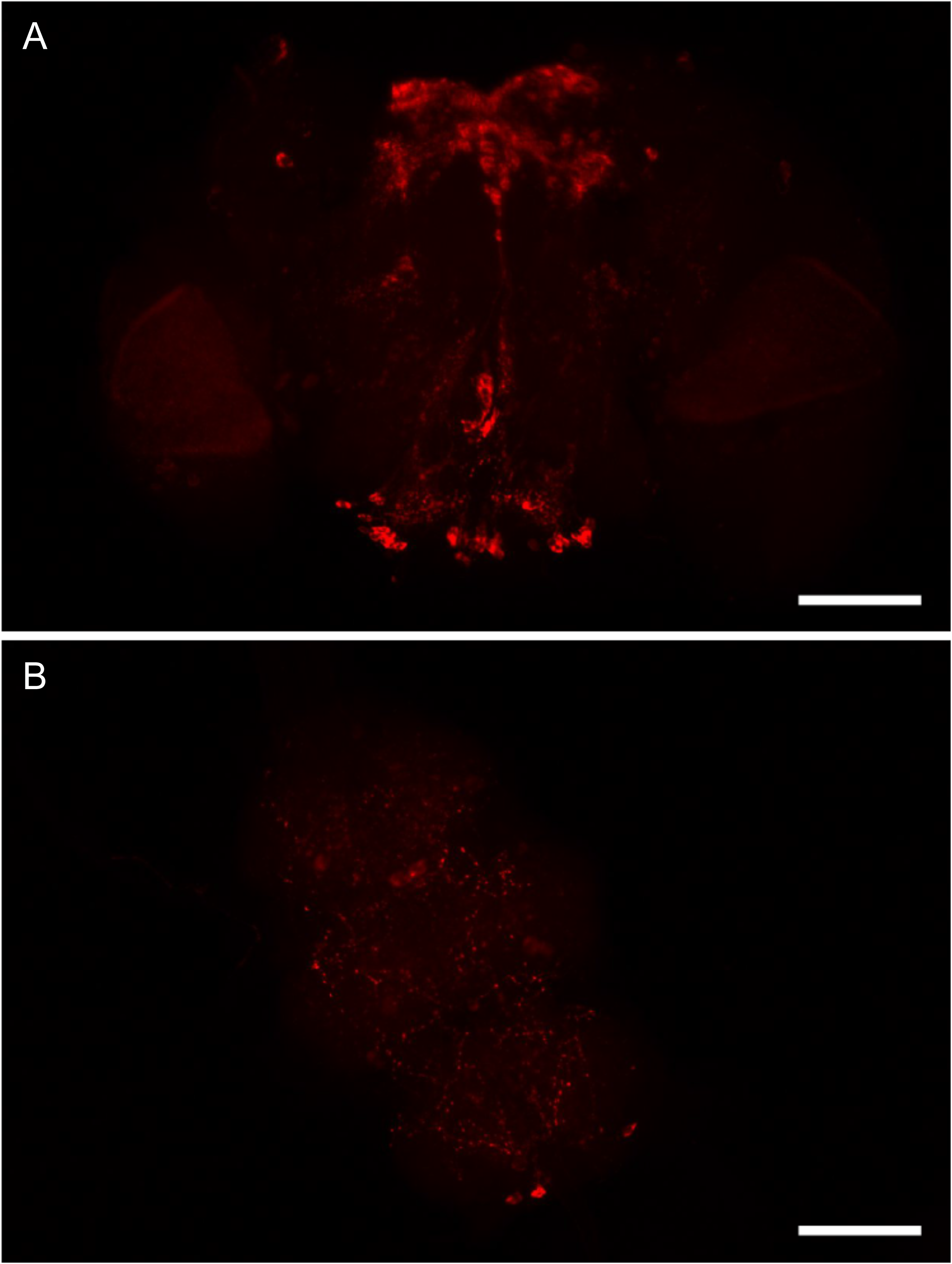
CAPA immunoreactivity observed in regions of the nervous system aside from the strongly-staining pair of neurosecretory cells in each of the abdominal ganglia. (A) CAPA immunoreactive staining in the brain showing a bilateral pair of neurons in each hemisphere of the brain and immunoreactive processes in the central margin with unknown origin. In the posterior suboesophageal ganglion, a number of small bilaterally-paired neurons (20-30 cells total) were detected. (B) In the fused thoracic ganglia, CAPA immunoreactive processes were observed on the ventral surface, with no consistently detected immunorective neurons. Although a qualitative observation, CAPA immunoreactive staining was substantially weaker in the brain, SOG and thoracic ganglia since exposure and gain settings on the fluorescence microscope were adjusted substantially to enable detection of weak immunoreactive staining. Scale bars: 100µm.

**Figure S4.**
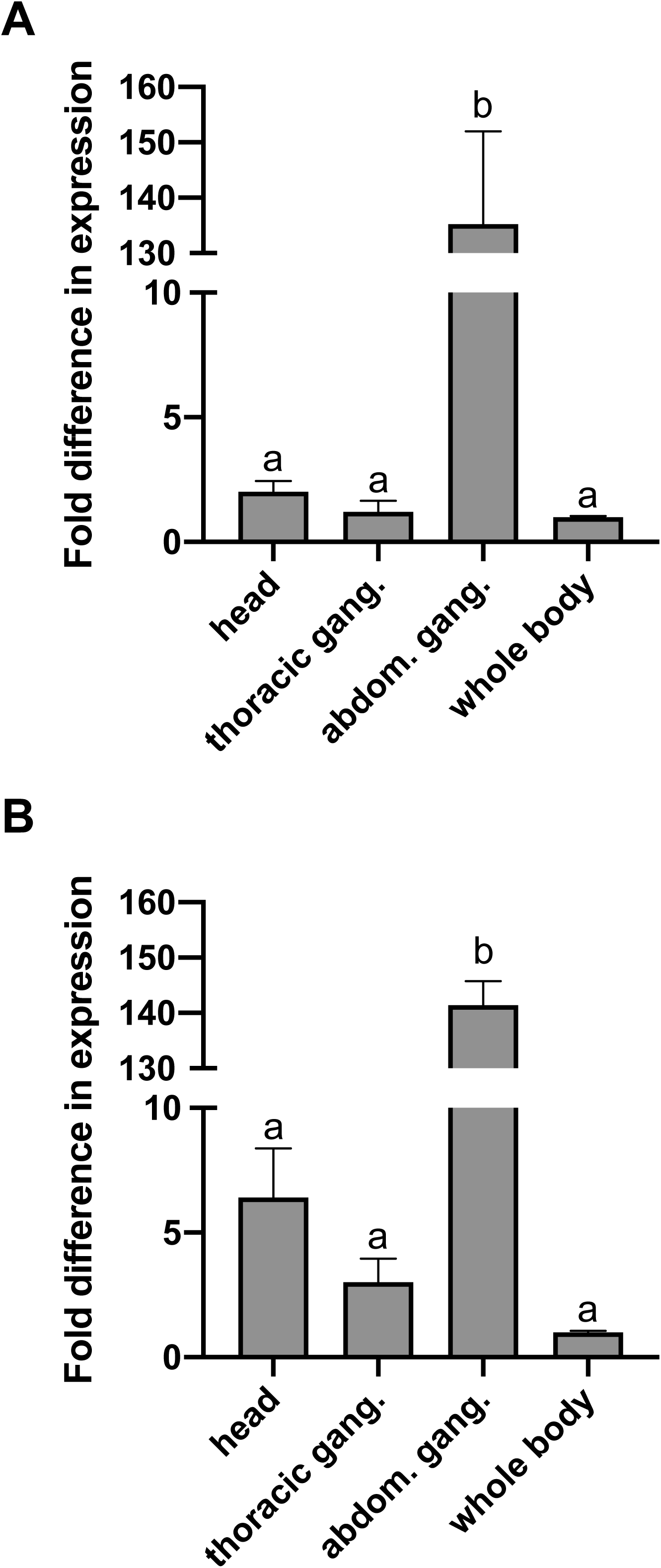
Expression analysis of CAPA neuropeptide (anti-diuretic hormone) transcript in different regions of the nervous system relative to whole adult (A) male and (B) female *A. aegypti* mosquitoes. Different letters denote bars that are significantly different from one another as determined by one-way ANOVA and Tukey’s multiple comparison post-hoc test (p < 0.01). Data represent the mean ± standard error (n = 3).

**Table S1.**
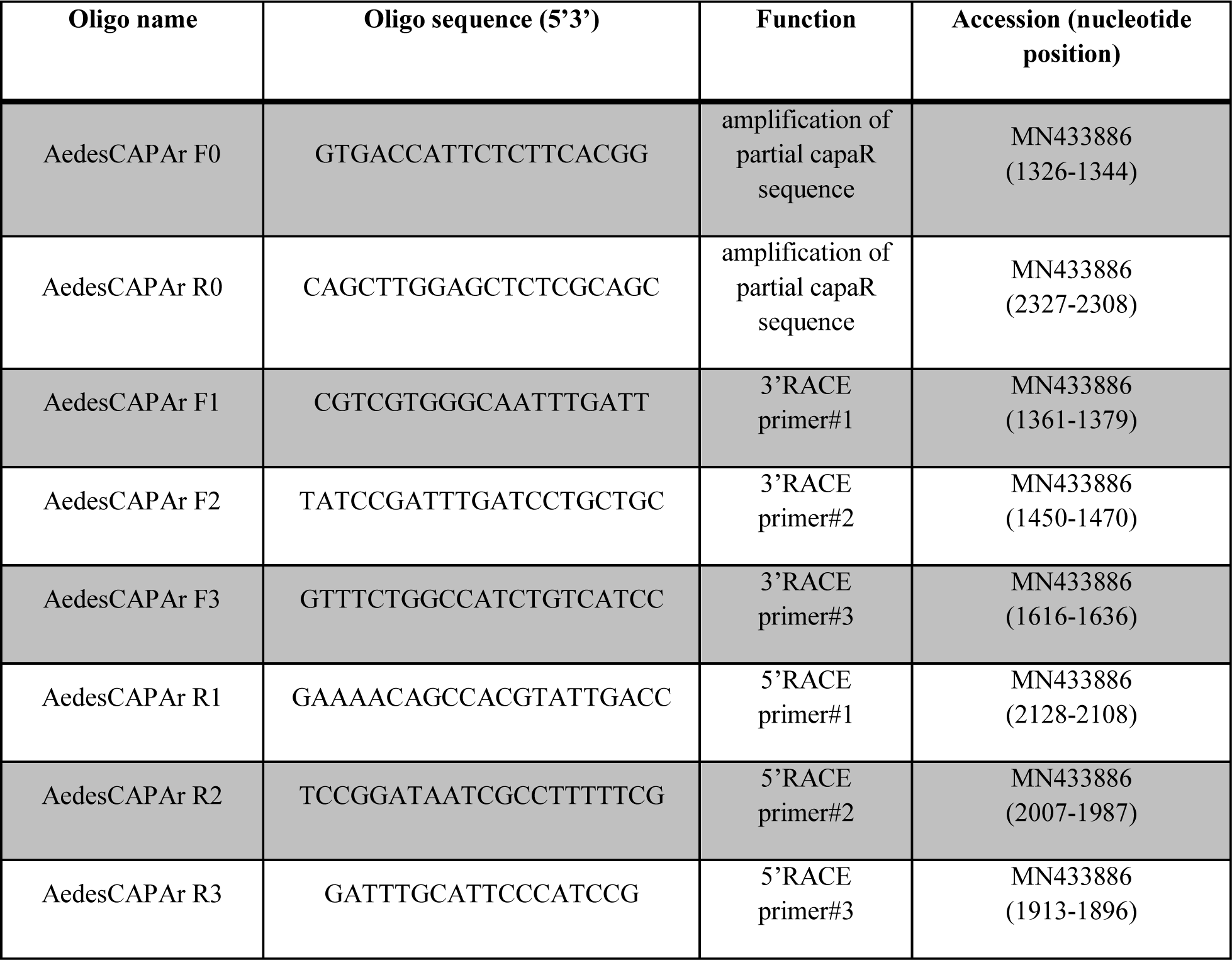
Oligonucleotides used for initial amplification and subsequent identification of the complete cDNA (including 5’ and 3’ UTR) encoding the *Aedes aegypti* anti-diuretic hormone (CAPA) receptor.

**Table S2.**
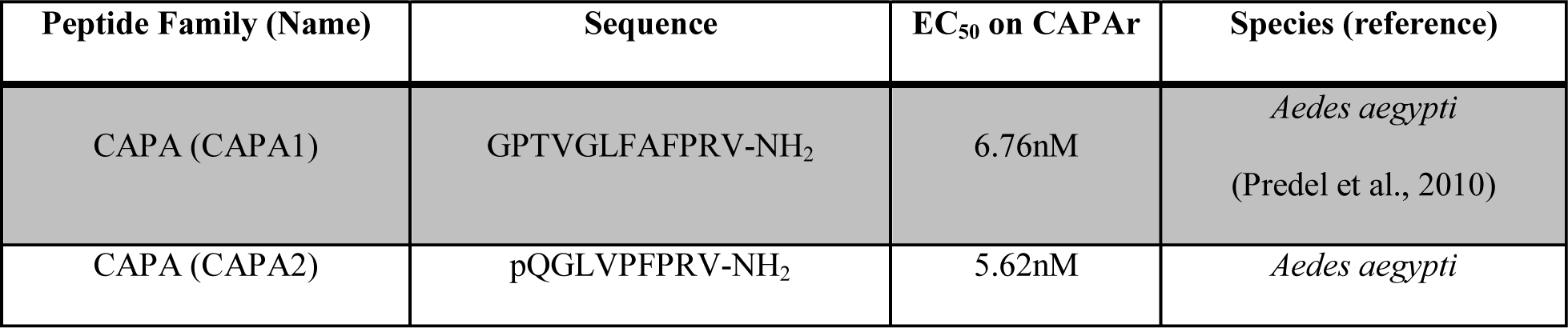
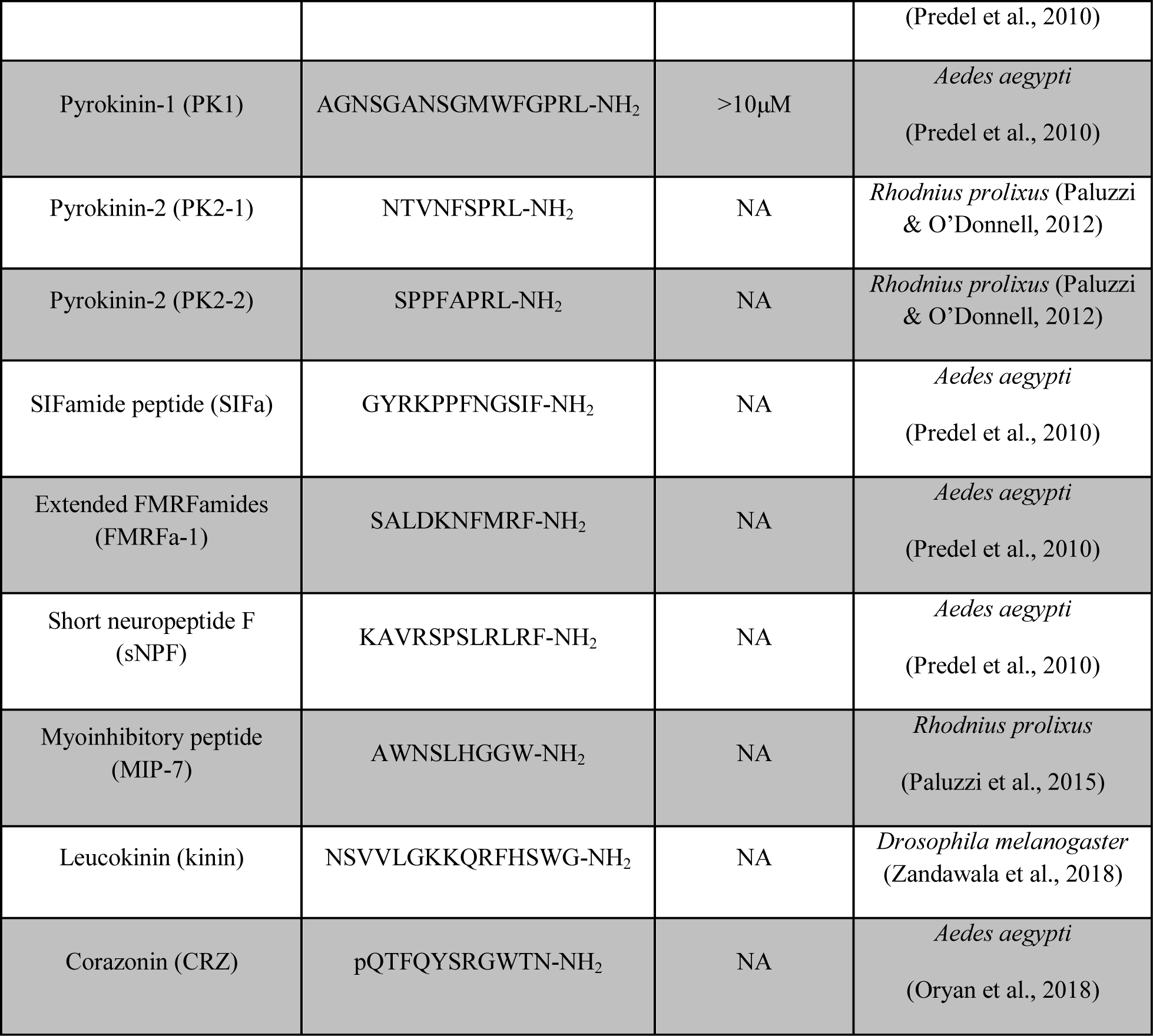
List and primary structure of several insect neuropeptides tested for functional activation of the mosquito anti-diuretic hormone (CAPA) receptor using heterologous bioassay. NA denotes peptides with no detectable activity when tested up to 10μM.

**Table S3.**
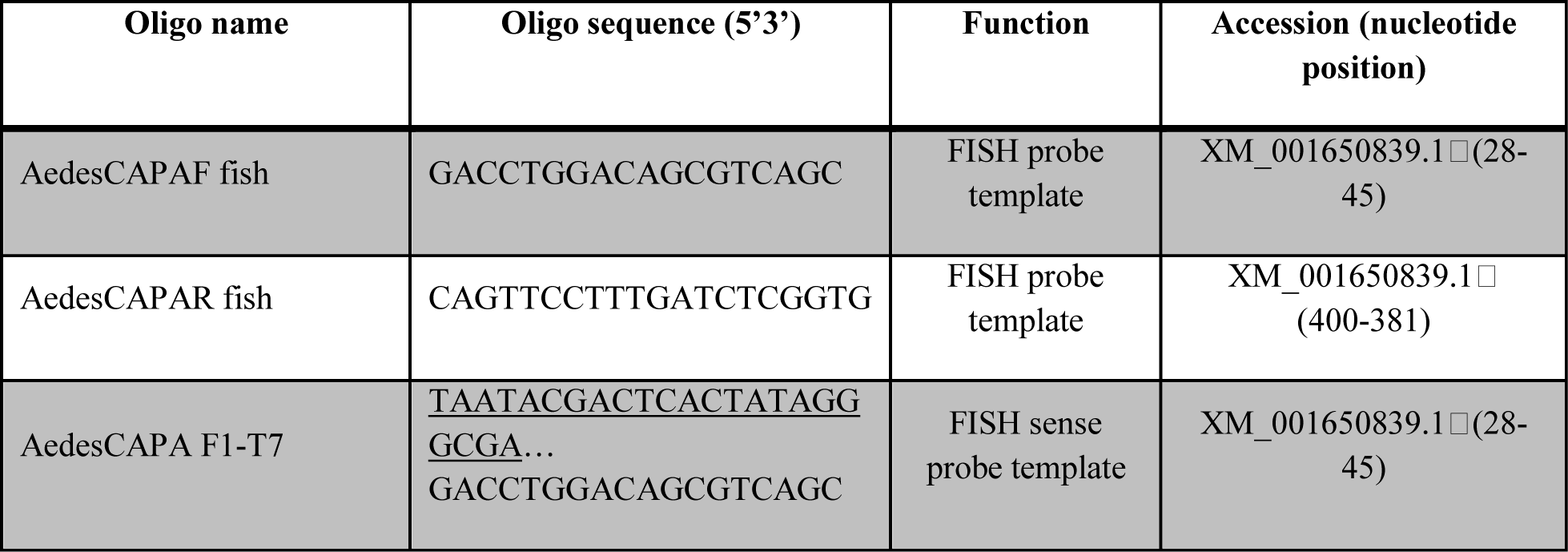
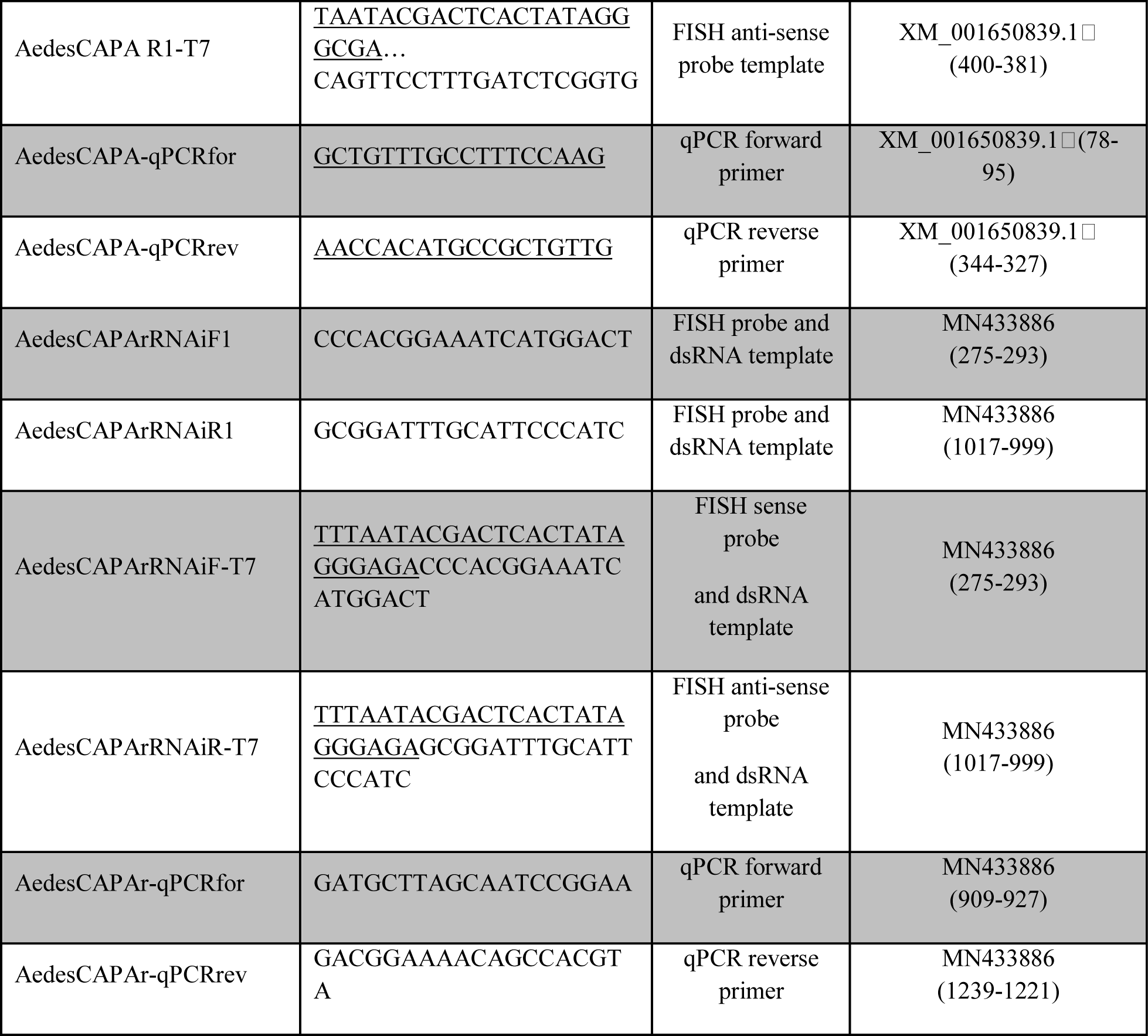
Oligonucleotides used for generation of fluorescent *in situ* hybridization probes, templates for *in vitro* dsRNA synthesis and gene-specific primers for quantitative PCR of the *Aedes aegypti* anti-diuretic hormone (CAPA) receptor.

